# Adaptive sampling–based enrichment enables genome reconstruction of intracellular symbionts despite host background and reference divergence

**DOI:** 10.64898/2026.03.25.714109

**Authors:** Wei-Kang Huang, Cheng-Hao Yang, Hui Chung, Yen-Chen Lee, Yung-Chieh Wu, Yen-Tong Chen, Mei-Hsiu Wan, Wen-Sheng Yeh, Yu-Ping Hong, Tsung-Hua Wu, Jian-Chiuan Li, Wei-Liang Liu, Chun-Hong Chen, Ying-Tsong Chen

## Abstract

Recovering genomes of intracellular microbes from host-dominated samples remains a major challenge in microbial genomics, due to low target abundance, overwhelming host DNA, and the inability to culture these organisms independently. Despite extensive interest in *Wolbachia*, efficient genome recovery directly from host tissues remains limited by the inefficiency of host-dominated sequencing and the constraints of existing enrichment strategies. Here, we demonstrate that Oxford Nanopore adaptive sampling (AS) enables efficient, real-time enrichment of target DNA directly from complex host tissues, providing a culture-free approach for genome recovery in such systems. To our knowledge, this represents the first application of enrichment-mode adaptive sampling to achieve de novo reconstruction of an intracellular endosymbiont genome in a mosquito system. Using *Aedes aegypti* mosquitoes infected with a locally derived *w*AlbB-like strain, we applied enrichment-mode AS to selectively sequence Wolbachia DNA. This resulted in an increase from <1% *Wolbachia* reads in conventional shotgun data to ∼90% under adaptive sampling. De novo assembly of AS-enriched long reads yielded a near-complete genome (∼1.5 Mb) in two contigs with >96–99% completeness. Comparative analyses revealed multiple large-scale chromosomal rearrangements relative to the reference *w*AlbB genome, demonstrating that adaptive sampling does not impose reference-dependent genome structure. Annotation further identified three prophage-associated regions, including two strain-specific expansions absent from the reference genome. Notably, cytoplasmic incompatibility genes (*cifA* and *cifB*) were identified adjacent to one of these regions, consistent with their known genomic association with prophage elements. Importantly, adaptive sampling remained effective despite substantial structural divergence between the reference and target genomes, revealing an unexpectedly robust application of this approach beyond its presumed operating conditions. Together, these results establish enrichment-mode adaptive sampling as a robust and scalable strategy for genome-resolved analysis of intracellular bacteria in host-associated systems.

## Introduction

Long-read sequencing technologies have greatly advanced microbial genomics by enabling contiguous, near-complete assemblies through shotgun sequencing. Despite these advances, recovering genomes from host-associated and intracellular microbes remains a major challenge, as target DNA is embedded within overwhelming amounts of host material, often at low abundance and without the possibility of independent cultivation. Under these conditions, conventional shotgun sequencing strategies are fundamentally limited in their ability to efficiently recover complete genomes, particularly in approaches that are not readily scalable across samples or study systems.

*Wolbachia* is a genus of intracellular α-proteobacteria that infects a wide range of arthropods and some nematodes (Hilgenboecker et al., 2008). These symbionts are of considerable interest due to their influence on host reproduction, physiology, and immunity, as well as their growing application in mosquito-borne disease control (Sinkins, 2004, Walker et al., 2011, Chouin-Carneiro et al., 2016). Many of these phenotypes, including cytoplasmic incompatibility (CI), are associated with genes encoded on prophage elements, which are frequently mobile and structurally variable within the *Wolbachia* genome. However, *Wolbachia* genomes are notoriously difficult to sequence because the bacteria cannot be cultured outside host cells and are often present at low abundance in specific tissues. Because *Wolbachia* genomes (∼1.5 Mb) are several orders of magnitude smaller than those of their mosquito hosts (∼1.4 Gb), this results in a pronounced imbalance in sequencing representation, where symbiont-derived reads are greatly outnumbered by host DNA even when the bacteria are present within infected tissues. Historically, most *Wolbachia* genome sequencing efforts have relied on short-read platforms such as Illumina to obtain the ultra-deep coverage needed to detect *Wolbachia* reads within host DNA. While effective, this strategy often produces highly fragmented assemblies, particularly in regions rich in insertion sequences, ankyrin-repeat genes, and other repetitive or mobile elements, which represent a pervasive feature of *Wolbachia* genomes and have long complicated shotgun-based assembly (Wu et al., 2004; Kaur et al., 2017; Siozios et al., 2013).

Several alternative strategies have been used to address these challenges. One approach involves maintaining *Wolbachia* in permissive insect cell lines and enriching bacterial DNA prior to sequencing, which has been used to recover *Wolbachia* genomic sequences and to produce a complete *w*AlbB genome from cell-line material (Mavingui et al., 2005; Sinha et al., 2019; Neupane et al., 2022). Another approach applies physical enrichment directly to infected host tissues to reduce host DNA contamination before sequencing, as demonstrated in *Aedes aegypti* using density-gradient centrifugation and other fractionation steps (Martinez et al., 2022). Although effective for obtaining complete *Wolbachia* genomes, this method requires multiple pre-sequencing manipulations, specialized laboratory infrastructure, and its enrichment efficiency can vary substantially, particularly in samples with low infection levels or field-derived material. A third approach mines *Wolbachia* reads in silico from publicly available host shotgun datasets and assembles them into draft genomes (Scholz et al., 2020). Collectively, these approaches have generated valuable references but remain labor-intensive, resource-dependent, or impractical for large-scale ecological or epidemiological studies involving diverse or field-collected samples.

Adaptive sampling is a software-defined real-time targeting feature in Oxford Nanopore sequencing that enables users to enrich specific DNA molecules during sequencing without prior amplification (Martin et al, 2022). As each DNA strand begins to pass through the nanopore, the signal generated by the first few hundred bases is basecalled and aligned in real time against a reference sequence. Based on this on-the-fly mapping, the system decides whether to continue sequencing the strand or to eject it from the pore. Earlier targeted sequencing methods, such as PCR-based enrichment, introduced amplification bias, required custom primers, and lacked flexibility. In contrast, adaptive sampling achieves sequence selectivity purely through software configuration, representing a major conceptual shift in how targeted sequencing can be performed. The platform currently supports two modes: a depletion mode that rejects reads mapping to a reference and an enrichment mode that retains those matching a given target. Regardless of mode, adaptive sampling has been applied in two broad contexts: (i) targeted sequencing of specific genomic loci and (ii) selective sequencing within complex metagenomic samples.

In the depletion mode, one study used adaptive sampling to sequence whole aphid samples containing their intracellular symbionts while depleting host reads in real time, enabling the direct recovery of complete *Wolbachia* and *Buchnera* genomes. This work showed that selective rejection of host DNA can substantially increase the representation of endosymbionts in the sequencing output, particularly when the host genome is relatively small and the symbiont is present at high abundance (Badger et al., 2024). However, applying this approach to more complex systems, such as mosquitoes with larger genomes and typically lower *Wolbachia* titers, may be less effective. In addition, depletion-mode strategies require high-quality host reference genomes, which are not always available for diverse or poorly characterized species.

By contrast, in the enrichment mode, adaptive sampling has been applied to targeted sequencing of specific genomic loci, such as human short tandem repeat expansions (Stevanovski et al., 2022), and to selective sequencing within complex metagenomic samples, including pathogen detection in tick microbiomes (Kipp et al., 2023). It has also been used to recover low-abundance eukaryotic pathogens from host-rich samples, such as *Plasmodium falciparum* from human blood (De Meulenaere et al., 2024). These studies illustrate the versatility of enrichment-mode adaptive sampling for capturing diverse targets without PCR-based preprocessing.

Collectively, these depletion- and enrichment-mode applications highlight a growing application space for adaptive sampling beyond classical target sequencing, emphasizing its role in enabling genome recovery from complex, unprocessed biological samples. However, whether enrichment-mode adaptive sampling can be effectively applied to recover genomes of low-abundance, intracellular bacteria from complex host tissues remains unclear, particularly under conditions of high host background and reference divergence.

Building on this emerging direction, we applied enrichment-mode adaptive sampling to selectively sequence the genome of *Wolbachia* from *Ae. aegypti* mosquitoes that had been stably transinfected with *w*AlbB derived from Taiwanese *Aedes albopictus*. These mosquitoes were developed and validated by our collaborators at the National Health Research Institutes in Taiwan as part of a broader vector control initiative.

Using a publicly available *w*AlbB genome as the reference, we performed Nanopore adaptive sampling on total DNA extracted from whole mosquitoes and achieved sufficient enrichment of *Wolbachia* reads to support high-quality de novo assembly. The resulting draft genome spanned two contigs covering >99% of the reference and included multiple prophage insertions that differed in both gene content and chromosomal position.

These results demonstrate that enrichment-mode adaptive sampling can enable near-complete genome reconstruction of uncultivable intracellular bacteria directly from host-dominated samples, even in the presence of substantial host background and moderate reference divergence. To our knowledge, this represents the first successful application of enrichment-mode adaptive sampling to recover a *Wolbachia* genome from mosquitoes—a biologically challenging system where standard sequencing approaches are often ineffective due to high host background and low symbiont abundance.

## Results

### Adaptive sampling markedly enriches *Wolbachia* reads relative to standard shotgun sequencing

To compare the performance of adaptive sampling with conventional shotgun sequencing, we prepared a Nanopore ligation library from a transinfected Taiwanese *Ae. aegypti* line carrying *Wolbachia w*AlbB originally derived from local *Ae. albopictus*. The presence of *w*AlbB in this mosquito line had been independently confirmed by qPCR. Using a single R10.4 flow cell, half of the active pores were assigned to adaptive sampling in enrichment mode (AS, targeting the *Ae. albopictus w*AlbB reference, GenBank CP031221.1), while the remaining pores performed standard shotgun sequencing (non-AS). Initial MinKNOW output indicated that adaptive sampling mode generated a substantial number of short non-target reads with a median length of ∼500 bp, consistent with the ∼200–400 bp decision-window of MinKNOW that limits rejection of shorter DNA molecules (Fig. 1).

**Figure 1.**
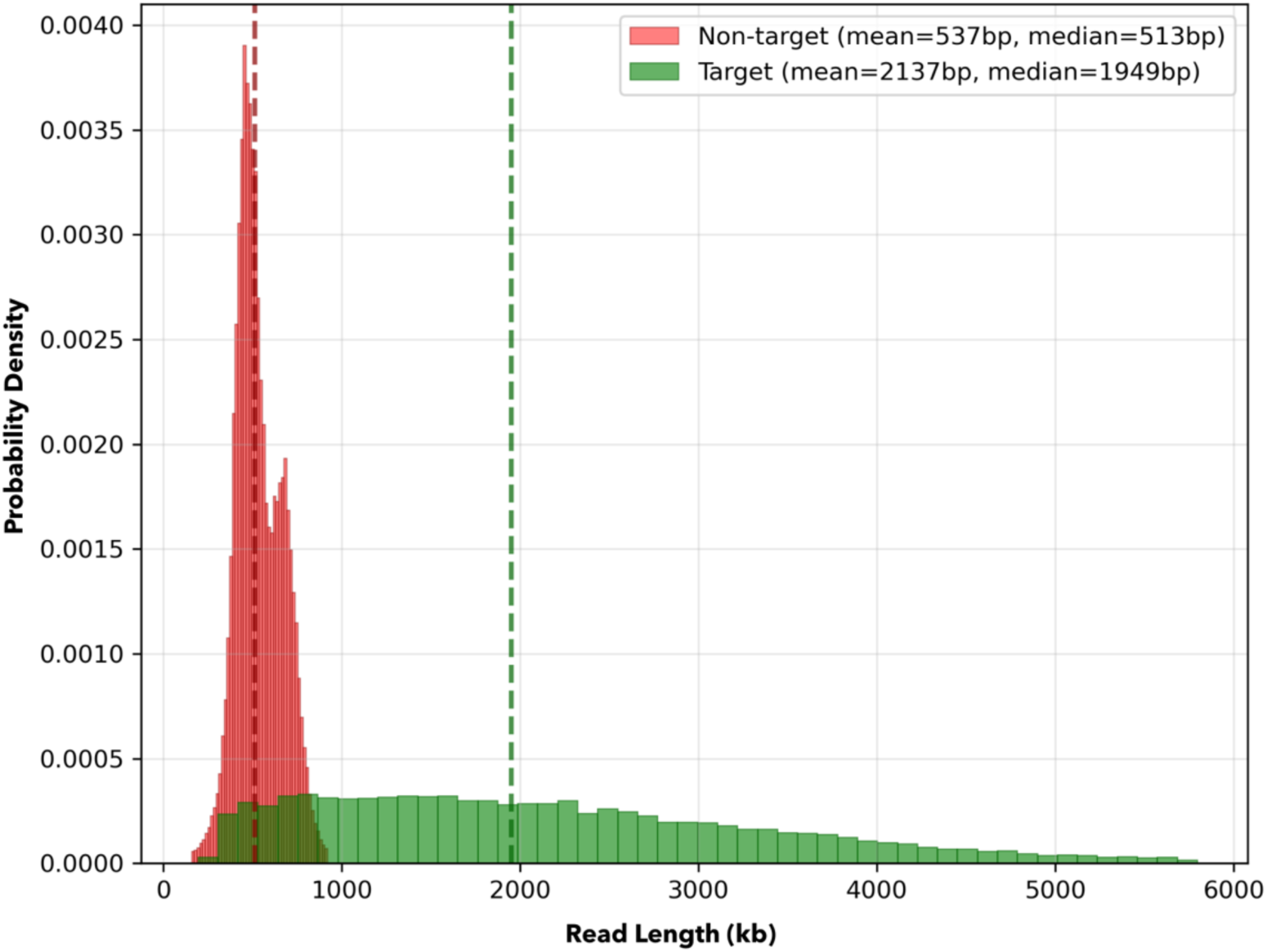
Read-length distributions reveal short non-target fragments retained during adaptive sampling. Non-target reads are strongly enriched in short fragments (mean ∼537 bp, median ∼513 bp), whereas target reads exhibit substantially longer lengths (mean ∼2137 bp, median ∼1949 bp). The accumulation of short non-target reads is consistent with the decision-window limitation of adaptive sampling, in which DNA molecules shorter than ∼200–400 bp cannot be reliably rejected during real-time classification. As a result, short non-target fragments are retained and sequenced, while longer on-target molecules are preferentially enriched. Outliers were removed for visualization.

To enable an accurate comparison, all reads generated within the same sequencing run (partitioned into AS and non-AS pores) were re-basecalled using Dorado in super-high-accuracy mode, and sequences shorter than 1 kb were removed. Taxonomic classification using Kraken2 revealed clear differences in the distribution of total sequenced bases between the two modes. In the shotgun dataset, approximately 70% of total bases were assigned to *Aedes* host sequences, whereas *Wolbachia* accounted for less than 1%. In contrast, the AS dataset was dominated by *Wolbachia*-derived sequences, which comprised nearly 90% of the total bases, representing more than a two-order-of-magnitude increase in *Wolbachia* sequence yield compared with shotgun sequencing (Fig. 2). Together, these results demonstrate that adaptive sampling substantially increases the representation of *Wolbachia* sequences in host-dominated samples, despite the large genome size disparity between host and symbiont.

**Figure 2.**
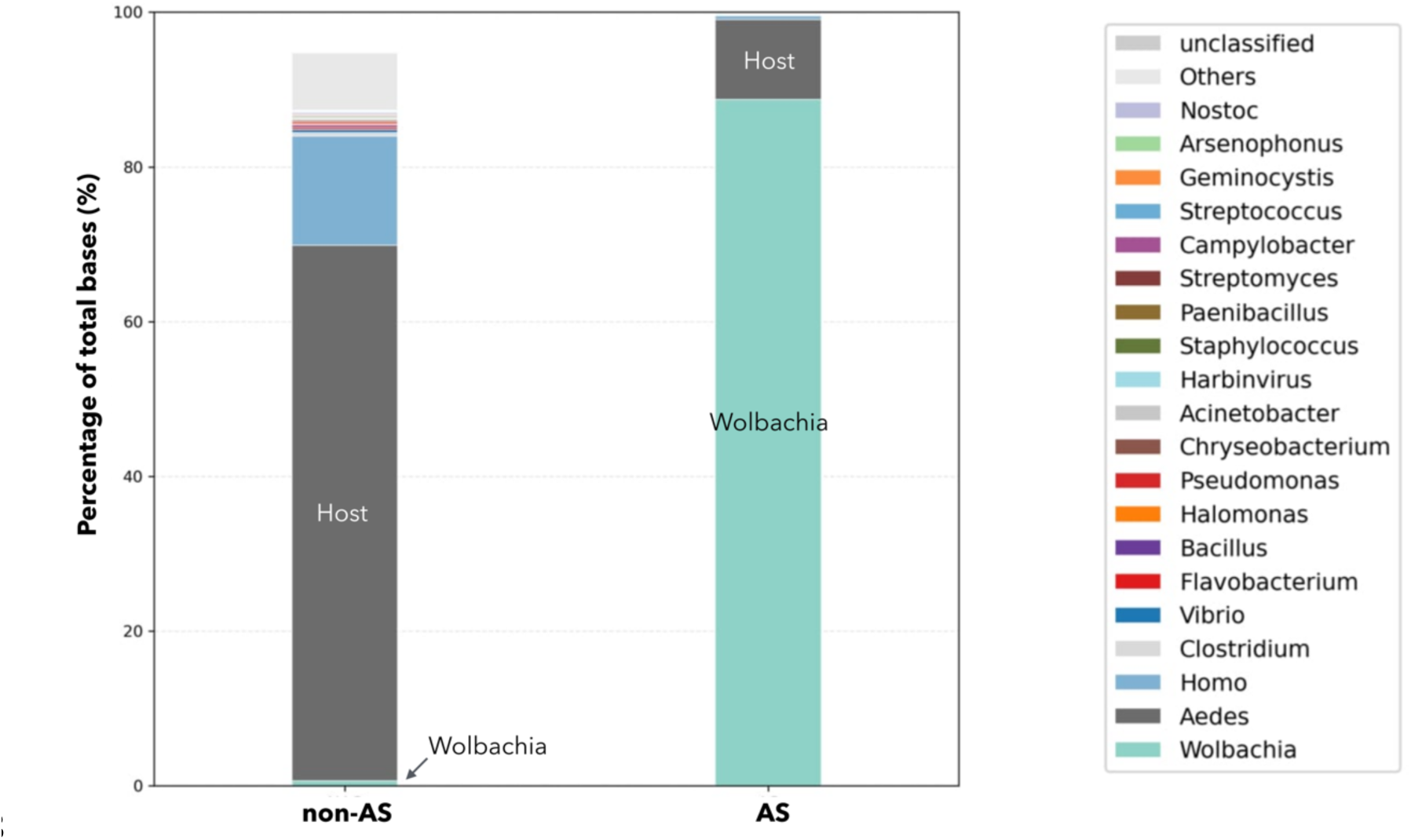
Adaptive sampling shifts taxonomic composition towards Wolbachia compared with conventional shotgun sequencing. Taxonomic composition of sequencing data generated by conventional shotgun sequencing (non-AS) and adaptive sampling (AS). Both datasets were derived from the same Nanopore ligation library prepared from a single DNA extraction of mosquito samples, ensuring identical input material. Sequencing was performed on a single flow cell using MinKNOW, with adaptive sampling and conventional sequencing conducted in parallel. Reads were classified at the genus level using Kraken2, and the proportion of total sequenced bases assigned to each taxon is shown. In the shotgun dataset (non-AS), host-derived sequences dominate, whereas adaptive sampling (AS) results in a marked increase in the proportion of *Wolbachia* sequences and a corresponding reduction in host-derived reads.

### Long-read filtering enables high-contiguity *Wolbachia* assembly and detects extensive chromosomal rearrangements

To evaluate whether the enriched long-read dataset was sufficient for *de novo* assembly, we analyzed sequencing output generated from a dedicated adaptive sampling run on a full R10.4 flow cell. We first characterized the sequencing output produced by adaptive sampling. A total of 6.8 Gb of ONT data were generated from the transinfected *Ae. aegypti* sample, comprising 2,917,865 reads with a mean length of 1,012 bp and an N50 of 2,950 bp. Because adaptive sampling retains a substantial number of short non-target reads, all sequences <5 kb were removed prior to assembly. This filtering step yielded 84,415 reads with an average length of 6,039 bp and an N50 of 5,891 bp, corresponding to >339× estimated coverage of the *Wolbachia* genome.

*De novo* assembly of the filtered dataset using Flye with default parameters readily produced two major contigs totaling ∼1.5 Mb, consistent with reported sizes of *w*AlbB-like *Wolbachia* strains. QUAST evaluation showed 96–97% genome fraction relative to the published reference (GenBank CP031221.1), ∼1% duplication, and no missing reference genes.

Whole-genome dot-plot alignment of the assembled contigs against the *Ae. albopictus w*AlbB reference genome (GenBank CP031221.1) revealed multiple forward and reverse diagonal blocks, indicating extensive chromosomal rearrangements between the assembled genome and the reference (Fig. 3). A prominent inversion was observed at the beginning of contig 1, while both contigs displayed non-collinear alignment patterns relative to the reference.

**Figure 3.**
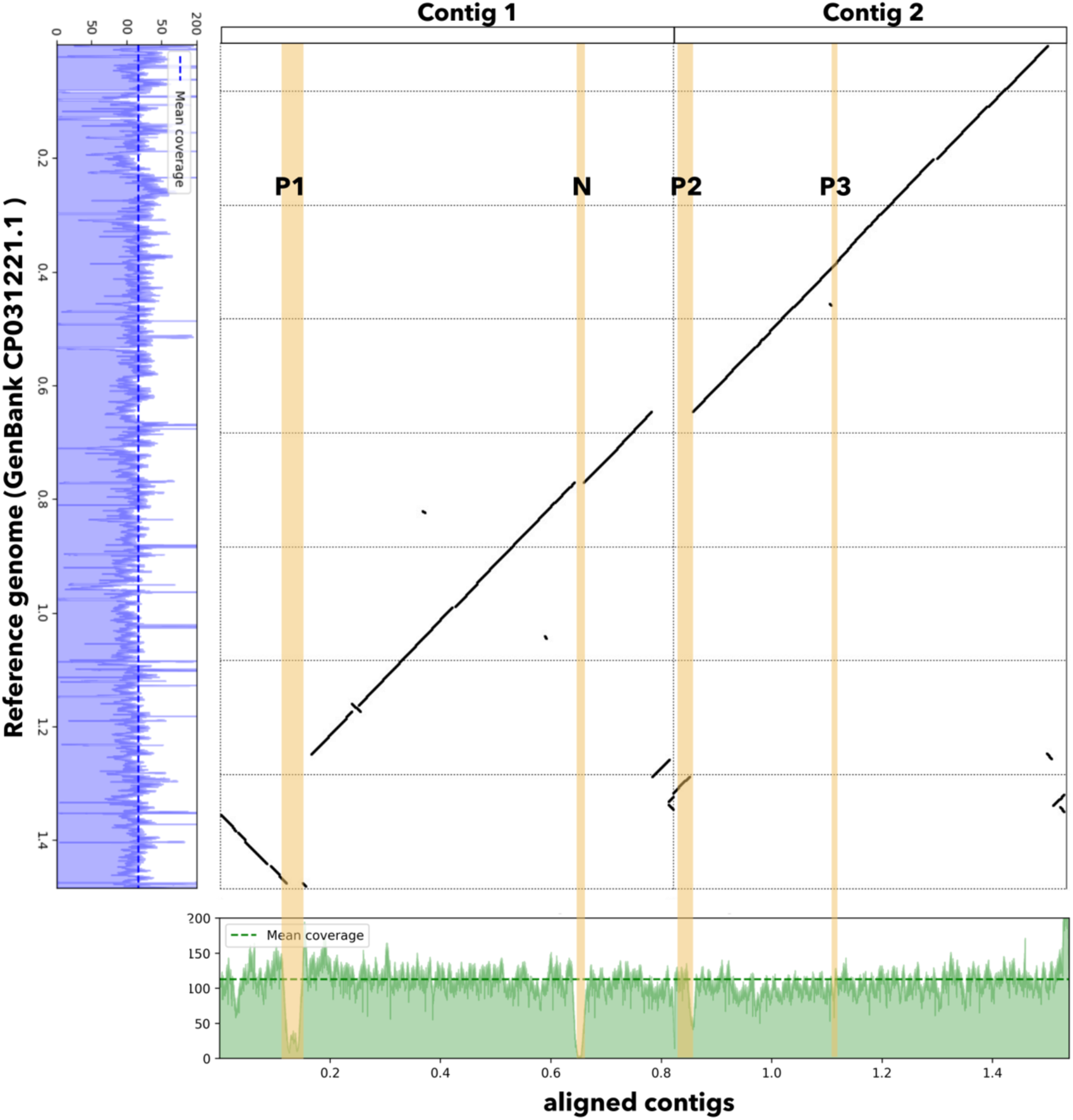
Whole-genome dot-plot alignment between the adaptive sampling-derived *Wolbachia* assembly and the *Ae. albopictus w*AlbB reference genome. The x-axis represents the two assembled contigs (contig 1 and contig 2, shown in order), and the y-axis represents the reference genome. Diagonal patterns with positive and negative slopes, as well as displaced alignment blocks, are observed in the dot plot, indicating large-scale chromosomal rearrangements between the assembled genome and the reference. Prophage-related regions (P1, P2, and P3) and a distinct genomic region absent from the reference genome (N) are annotated on the assembled contigs and highlighted by light yellow bands. Coverage profiles of adaptive sampling reads mapped to the reference genome (blue, left side) and to the assembled contigs (green, bottom) were generated using minimap2. Median coverage depths are approximately 110–120× for both mappings, with two broad low-coverage regions observed on the assembled contigs that correspond to P1 and region N.

Three prophage-related regions were identified on the assembled contigs and designated P1, P2, and P3 based on genome annotation. Regions P2 and P3 corresponded to prophage-associated loci present in the reference genome, whereas P1 was absent from the reference. In addition, a distinct genomic segment not present in the reference genome was identified and designated as region N. Coverage profiles of adaptive sampling reads mapped to the reference genome showed a relatively uniform depth across most regions, with a median coverage of approximately 110–120×. When the same reads were mapped to the assembled contigs, two broad regions of markedly reduced coverage (approximately 4–6×) were observed. These low-coverage regions coincided with P1 and region N, both of which were absent from the reference genome. These results suggest that adaptive sampling–derived reads extend beyond reference-matching regions and enable recovery of structurally divergent genomic segments, although coverage in these regions is reduced.

### Moderate long-read coverage is sufficient for near-complete *Wolbachia* genome recovery

Given the enrichment achieved by adaptive sampling, we next asked what level of sequencing depth is required for near-complete genome recovery. We performed a subsampling analysis using the filtered long-read dataset. Reads <5 kb were excluded, and the remaining reads were mapped to the reference using minimap2. Only reference-mapping reads were retained to construct a *Wolbachia*-enriched dataset representing the full sequencing depth. This dataset was randomly partitioned into subsets corresponding to 1–100% of the original data, and each subset was assembled de novo using Flye and evaluated with QUAST.

Assembly completeness increased sharply up to approximately 20× coverage and plateaued at moderate depth. Above ∼50× coverage, assemblies consistently recovered >95% of reference genes and exhibited stable assembly metrics. These results indicate that near-complete *Wolbachia* genome recovery can be achieved without excessive sequencing and provide a practical benchmark for determining when adaptive sampling runs can be terminated to optimize sequencing efficiency and cost-effectiveness.

### Long-read adaptive sampling captured three prophage-associated regions, including two strain-specific expansions

Genome annotation using NCBI PGAP, combined with prophage prediction using PHASTEST, identified three prophage-associated regions in the assembled *Wolbachia* genome. Comparative analysis with the *Ae. albopictus w*AlbB reference revealed that two of these regions contained additional sequence blocks not present in the reference genome—each fully resolved by long-read adaptive sampling data— suggesting strain-specific expansion or insertion-like variation (Fig. 4).

**Figure 4.**
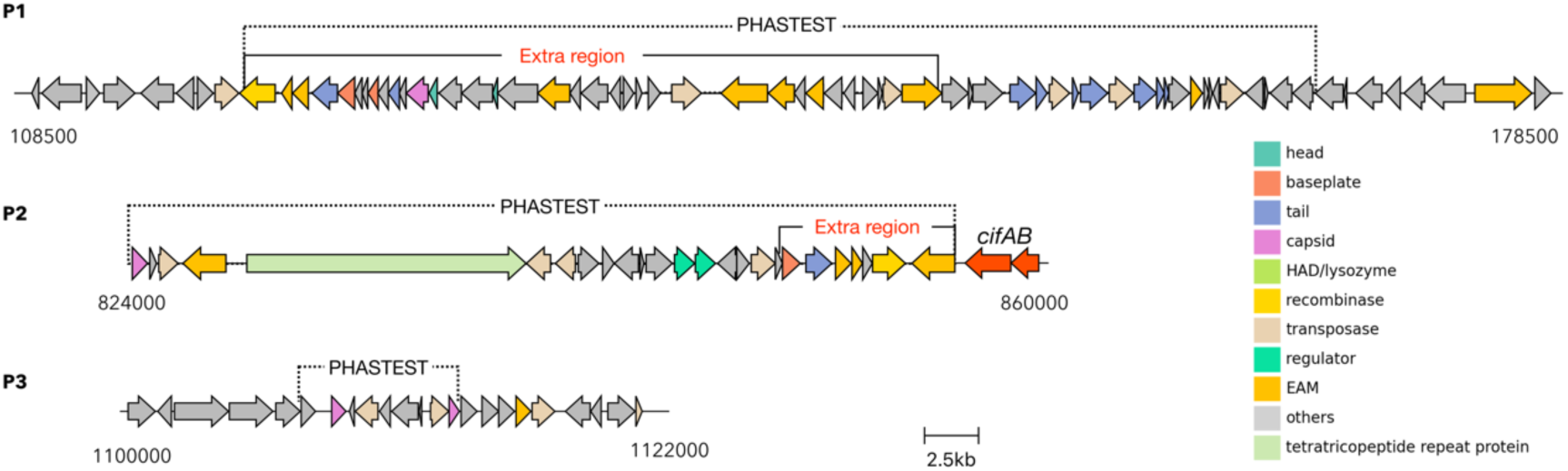
Annotation and comparative analysis of prophage-associated regions in the assembled *Wolbachia* contigs. Prophage-associated regions identified in the assembled *Wolbachia* contigs based on annotation with NCBI PGAP. Three regions (P1, P2, and P3) are shown across the two major assembled contigs. Arrows represent predicted open reading frames (ORFs), colored according to functional categories. Prophage regions predicted by PHASTEST are indicated by dashed boxes. Comparative analysis with the *Ae. albopictus w*AlbB reference genome (GenBank CP031221.1) revealed additional sequence segments (labeled as “extra region”) in P1 and P2 that are absent from the reference genome. Homologs of the cytoplasmic incompatibility genes *cifA* and *cifB* are highlighted in red and located adjacent to P2.

Prophage 1, approximately 50 kb in size, exhibited gene content characteristic of WO phages known to mediate cytoplasmic incompatibility (CI), including numerous genes belonging to the Eukaryotic Association Module (EAM).

Prophage 2, roughly half the size of Prophage 1, also contained an extra region absent from the *w*AlbB reference. At the terminal region adjacent to Prophage 2, we identified intact homologs of the CI-associated genes *cifA* and *cifB*, consistent with the known genomic association of CI loci with prophage-related elements in *Wolbachia*.

In contrast, Prophage 3 was structurally conserved and closely matched the corresponding prophage region in the reference genome.

Taken together, these findings indicate that long-read adaptive sampling enables near-complete *Wolbachia* genome reconstruction and resolves prophage-associated structural variation, including regions harboring CI-associated genes, and remains effective for genome reconstruction even when the target genome exhibits substantial structural divergence from the reference.

## Discussion

### Technical feasibility and innovation of the adaptive sampling approach

Recovering genomes of intracellular symbionts directly from host-dominated samples remains a fundamental challenge in microbial genomics, due to the combined constraints of low target abundance, overwhelming host DNA, and the inability to culture these organisms independently. This study demonstrates that adaptive sampling combined with Nanopore long-read sequencing enables near-complete recovery of a *Wolbachia* genome directly from whole mosquito DNA extracts— without the need for cell culture, bacterial isolation, or targeted amplification. Although adaptive sampling has been increasingly used in clinical metagenomics and human genome studies, our work represents the first successful application of enrichment-mode adaptive sampling to reconstruct an intracellular endosymbiont genome from host tissues. By addressing these constraints simultaneously, our results highlight the potential of this approach to overcome key limitations in sequencing host-associated microbes. This establishes a practical and accessible framework for genome-resolved analysis of intracellular symbionts that cannot be isolated in pure culture.

### Adaptive sampling retains short non-target fragments due to decision-window constraints

Although AS efficiently enriched *Wolbachia* DNA, we observed that a substantial subset of non-target reads remained in the enriched dataset. These sequences were almost exclusively short (<1 kb), a pattern consistent with the mechanistic constraints of adaptive sampling. Because MinKNOW requires ∼200–400 bp of sequence to make a keep/eject decision, DNA fragments shorter than this threshold traverse the pore before an eject command can be issued. Consequently, short host-derived fragments are inevitably retained.

These short reads disrupted assembly by introducing spurious edges in the assembly graph—an important consideration for *Wolbachia*, whose genomes contain repetitive insertion sequences, mobile elements, and structurally variable prophage regions. Removing reads <5 kb prior to assembly enabled the recovery of two long contigs (∼1.5 Mb) with clean structure, stable synteny, and no missing reference genes. This reveals an important methodological insight: adaptive sampling enriches the target by proportion, not by absolute exclusion, and size-based filtering is essential for high-quality genome assembly in host-dominated samples.

### Rationale and advantages of enrichment-mode adaptive sampling

Traditional approaches for *Wolbachia* genome sequencing—such as deep shotgun sequencing of host DNA or cell fractionation in insect lines—are inefficient and often insufficient for resolving repetitive regions like WO prophages. Many published Wolbachia genomes remain fragmented due to reliance on short-read sequencing.

In this study, we employed enrichment-mode adaptive sampling, using the *Ae. albopictus w*AlbB genome as a guide without requiring prior knowledge of the *Ae. aegypti* host genome. While exclusion-mode adaptive sampling could theoretically deplete host DNA, its effectiveness depends on the completeness and accuracy of the host reference. In mosquito-associated systems, host genome assemblies may be incomplete and can contain sequences of symbiont origin or regions with high similarity to *Wolbachia*, raising the risk of unintended exclusion of target reads. In addition, removal of host-derived sequences does not eliminate other sources of background DNA, such as microbiota or diet-derived material, which may still dominate the sequencing output.

Taken together, these considerations make enrichment-mode adaptive sampling a more reliable strategy for selectively capturing *Wolbachia* sequences in complex host-associated samples. This strategy successfully provided deep target coverage and enabled near-complete assembly, highlighting its applicability to endosymbionts residing in complex host tissues.

### Robust genome recovery and structural resolution despite reference divergence

A key advantage of enrichment-based adaptive sampling is that perfect identity between the reference and target genomes is not required. In this study, the reference genome used for adaptive sampling originated from a *w*AlbB strain infecting *Ae. albopictus*, whereas the sequenced bacterium represents a *w*AlbB-like strain experimentally transinfected into *Ae. aegypti*. Despite this divergence, adaptive sampling efficiently enriched *Wolbachia* DNA and enabled near-complete genome recovery.

Although adaptive sampling is reference-guided, our results demonstrate that enrichment is governed by sequence similarity rather than by preservation of reference collinearity. Long reads spanning rearranged regions were retained and incorporated into the final assembly, enabling resolution of extensive inversions and synteny disruptions relative to the reference genome. Dot-plot comparisons revealed mosaic patterns of forward and reverse alignment blocks, consistent with large-scale chromosomal rearrangements previously reported in *Wolbachia* comparative genomics studies (Vostokova et al., 2023; Jacobs et al., 2024; Muro et al., 2023). These rearrangements were supported by long reads spanning junctions, supporting their authenticity and excluding assembly artifacts.

Importantly, adaptive sampling did not restrict genome reconstruction to reference-matching regions. Prophage-associated segments and distinct genomic regions absent from the reference genome were successfully recovered and assembled as contiguous sequences, despite reduced coverage relative to reference-aligned regions. Notably, such divergent regions are often associated with prophage-related elements. In our assembly, *cifA* and *cifB* genes were identified adjacent to one of the prophage-associated regions, consistent with previous reports linking cytoplasmic incompatibility loci to prophage elements in *Wolbachia*. Together, these findings demonstrate that adaptive sampling enables recovery of structurally variable genomic regions that extend beyond reference-matching sequences, even when the target genome exhibits substantial structural divergence from the reference.

Beyond its methodological implications, the extensive structural divergence observed in the Taiwanese transinfected *w*AlbB genome provides insight into the intrinsic plasticity of *Wolbachia* genome architecture. Accumulating evidence indicates that *Wolbachia* genomes are highly dynamic, with frequent rearrangements mediated by insertion sequences and prophage elements. The pronounced differences between the transinfected *Ae. aegypti w*AlbB-like strain and the published *Ae. albopictus* reference genome are consistent with this emerging view and suggest that genome restructuring is an ongoing process rather than a rare evolutionary event. Such plasticity may reflect both pre-existing geographic variation and adaptive responses following transinfection into a novel mosquito host, underscoring the evolutionary flexibility of *Wolbachia* in new host environments.

### Resolving prophage diversity and CI-associated elements with long-read sequencing

Wolbachia genomes frequently harbor WO prophages, which contribute to genomic plasticity and often encode cytoplasmic incompatibility factors. These elements are typically repetitive, mosaic, and difficult to resolve with short-read data. Our long-read assembly recovered three prophage-related regions, two of which contained expanded or insertion-like segments not present in the reference genome. Notably, one prophage region included intact *cifA* and *cifB* homologs, consistent with its potential role in CI.

The ability to resolve such regions illustrates the value of long-read sequencing paired with adaptive sampling. By capturing structural variation and mobile genetic content in their entirety, this approach enables deeper analysis of *Wolbachia*’s evolutionary dynamics, host-interaction mechanisms, and strain-specific functional variation.

### Toward scalable genome sequencing of intracellular symbionts

The strategy described here—adaptive sampling combined with long-read sequencing—provides a robust, scalable, and culture-free framework for recovering high-quality genomes of intracellular bacteria directly from host tissues. Our results show that near-complete *Wolbachia* assemblies can be obtained from complex mosquito samples, despite host contamination and reference divergence. This approach reduces dependency on host genomic resources, eliminates PCR enrichment biases, and provides sufficient long-read continuity to resolve repetitive and mobile elements.

More broadly, our findings indicate that enrichment-mode adaptive sampling can enable genome-resolved analysis of host-associated microbial systems, particularly for intracellular and uncultivable bacteria that are difficult to access using conventional sequencing approaches. By simultaneously tolerating host-dominated backgrounds and moderate reference divergence, this strategy expands the practical scope of genome recovery for intracellular and uncultivable bacteria.

With further optimization, it could be applied to field-collected specimens and genomic surveillance of *Wolbachia*-based vector control programs—particularly in settings where reference genomes may be incomplete or divergent. Together, our findings demonstrate that adaptive sampling enables robust and structurally resolved genome reconstruction in host-associated systems, even in the presence of substantial host contamination and reference divergence.

## Materials and Methods

### Mosquito strain, *Wolbachia* transinfection, and sample preparation

Mosquito tissues used in this study were obtained from a *Wolbachia*-transinfected *Ae. aegypti* line, designated *w*AlbB-Tw-Kao. This line was generated by embryonic microinjection of cytoplasm derived from field-collected *Ae. albopictus* mosquitoes in Kaohsiung, Taiwan, and has since been maintained at the National Mosquito-Borne Diseases Control Research Center, National Health Research Institutes (NHRI), Taiwan. The establishment and biological characterization of this transinfected line, including cytoplasmic incompatibility and viral interference phenotypes, are described elsewhere (Hui-Ying Yu et al., manuscript under revision).

### Genomic DNA extraction and quality assessment

Total genomic DNA was extracted from pooled *w*AlbB-Tw-Kao adult mosquitoes using the DNeasy Blood & Tissue Kit (Qiagen, Hilden, Germany), following the manufacturer’s protocol with minor modifications. Approximately 50 mg of mosquito tissue was used per extraction, corresponding to roughly twice the recommended input amount. Prior to enzymatic lysis, tissue samples were briefly subjected to sonication to enhance mechanical disruption and improve lysis efficiency. Following purification, DNA concentration was measured using a Qubit fluorometer (Thermo Fisher Scientific, Waltham, MA, USA).

### Nanopore library preparation and adaptive sampling sequencing

Sequencing libraries were prepared from total genomic DNA using the Oxford Nanopore Technologies (ONT) ligation sequencing kit (LSK114), following the manufacturer’s instructions. Prepared libraries were loaded onto an R10.4 flow cell and sequenced on a MinION Mk1B device (ONT, Oxford, UK).

Adaptive sampling was enabled in enrichment mode to selectively retain Wolbachia-derived reads during sequencing. The *w*AlbB reference genome from *Ae. albopictus* (GenBank accession no. CP031221.1) was specified as the target sequence for on-device enrichment. Sequencing runs were controlled using MinKNOW v24.11.8 (Core v6.2.6) on a computer workstation running Ubuntu 22.04 LTS. Raw nanopore signal data were initially basecalled in real time during sequencing and subsequently re-basecalled using Dorado v7.6.7 (ONT) in super high-accuracy (SUP) mode. All basecalling procedures were accelerated using an NVIDIA RTX4000 Ada GPU. Filtering of short reads was performed using filtlong (v0.2.1).

### Assessment of adaptive sampling enrichment efficiency

To evaluate the enrichment efficiency of adaptive sampling, a controlled sequencing experiment was performed using a single R10.4 flow cell. The same sequencing library was used to generate two datasets under different conditions: conventional shotgun sequencing with adaptive sampling disabled, followed by enrichment-mode adaptive sampling activated during the same run. Reads from the two phases were separated based on sequencing time and run configuration. All reads were basecalled in super high-accuracy (SUP) mode, and reads shorter than 1 kb were removed prior to downstream analysis. Taxonomic classification was performed using Kraken2 with the standard reference database. Taxonomic profiles derived from the conventional shotgun and adaptive sampling–enriched datasets were compared to assess changes in species-level composition resulting from adaptive sampling.

### Genome assembly, quality assessment, and annotation

De novo assembly of *Wolbachia* genome was performed using adaptive sampling– derived reads that were re-basecalled in super high-accuracy (SUP) mode and filtered to retain reads longer than 5 kb. Assemblies were generated using Flye (v2.9) with the --nano-hq preset optimized for high-quality nanopore reads. Assembly quality and contiguity were assessed using QUAST (v5.2.0), with standard assembly metrics including total assembly length, number of contigs, and N50. Genome completeness was further evaluated using BUSCO (v5.4.3) with the appropriate bacterial lineage dataset to assess the recovery of conserved single-copy genes. Structural and functional annotation of the assembled *Wolbachia* genome was performed using the NCBI Prokaryotic Genome Annotation Pipeline (PGAP) with default parameters. Prophage regions and phage-associated genes were identified using PHASTER (PHAge Search Tool Enhanced Release).

### Comparative genomics and structural analysis

Pairwise genome alignments between the assembled *Wolbachia* contigs and the Aedes albopictus *w*AlbB reference genome (GenBank accession no. CP031221.1) were performed using NUCmer from the MUMmer4 package. Genome-wide collinearity and structural concordance were visualized using dot-plot analyses based on the resulting alignments. Coverage depth of adaptive sampling–derived reads across the reference genome was calculated by read alignment and summarized to provide supporting evidence for assembly continuity.

## Data availability

The assembled Wolbachia wAlbB-Tw-Kao genome generated in this study has been deposited in GenBank under accession number JBWDTH000000000. The associated BioSample is available under accession number SAMN54601416 (sample name: Wolbachia_Tw-Kao).

Raw Nanopore sequencing reads have been deposited in the Sequence Read Archive (SRA) under accession numbers SRR37737917 (adaptive sampling and conventional sequencing performed in parallel on a single flow cell) and SRR37737918 (adaptive sampling run). All data are publicly available.

## Acknowledgements

We are grateful to the current and former members of our laboratory, many of whom contributed to this study alongside their own independent research projects. Their sustained interest, voluntary involvement, and continued engagement—even after graduation—provided essential motivation and intellectual support for this work.

